# Astrocyte subdomains respond independently in vivo

**DOI:** 10.1101/675769

**Authors:** Mónica López-Hidalgo, Vered Kellner, James Schummers

## Abstract

Astrocytes contact thousands of synapses throughout the territory covered by its fine bushy processes. Astrocytes respond to neuronal activity with an increase in calcium concentration that is in turn linked to their capacity to modulate neuronal activity. It remains unclear whether astrocytes behave as a single functional unit that integrates all of these inputs, or if multiple functional subdomains reside within an individual astrocyte. We utilized the topographic organization of ferret visual cortex to test whether local neuronal activity can elicit spatially restricted events within an individual astrocyte. We monitored calcium activity throughout the extent of astrocytes in ferret visual cortex while presenting visual stimuli that elicit coordinated neuronal activity spatially restricted to functional columns. We found visually-driven calcium responses throughout the entire astrocyte that was largely independent in individual subdomains, often responding to different visual stimulus orientations. A model of the spatial interaction of astrocytes and neuronal orientation maps recapitulated these measurements, consistent with the hypothesis that astrocyte subdomains integrate local neuronal activity. Together, these results suggest that astrocyte responses to neural circuit activity are dominated by functional subdomains that respond locally and independently to neuronal activity.

## Introduction

Astrocytes are a prominent non-neuronal cell type in the brain. They are intimately connected with neural circuit elements, and respond to neuronal activity with increases in cytosolic calcium concentration (Bazargani and Attwell, 2016; Cornell-Bell et al., 1990; Khakh and McCarthy, 2015; Rusakov, 2015; Shigetomi et al., 2016; Volterra et al., 2014). These calcium responses have been shown to be necessary for initiating signaling cascades that influence brain function, including metabolic support (Harris et al., 2012), regulation of synaptic transmission (Di Castro et al., 2011; Panatier et al., 2011) and synaptic plasticity (Perea and Araque, 2007). Astrocyte calcium activity has been shown to be elicited by a number of neurotransmitter systems, including both classic synaptic neurotransmitters (Di Castro et al., 2011; Panatier et al., 2011; Porter and McCarthy, 1996), and volume-transmitted neuromodulators (Chen et al., 2012; Ding et al., 2013; Lopez-Hidalgo et al., 2012; Papouin et al., 2017; Paukert et al., 2014; Salm and McCarthy, 1992; Takata et al., 2011). Astrocyte calcium signaling has been demonstrated to be necessary for normal neural circuit function (Oliveira et al., 2015), and to be involved in numerous pathological brain states (Barres, 2008; Chung et al., 2015; Verkhratsky and Parpura, 2016). Astrocytes have been implicated in modulating neuronal activity at spatial scales ranging from synchronizing large populations of neurons (Fellin et al., 2004; Pirttimaki et al., 2017; Poskanzer and Yuste, 2016), to regulating individual synapses (Di Castro et al., 2011; Panatier et al., 2011). Defining the spatial characteristics of astrocyte calcium signaling is therefore critical for understanding the role of astrocytes in brain function.

Cell-wide calcium events can be elicited *in vivo*, in response to sensory or motor-related neural activity, both under anesthetized and awake conditions (Paukert et al., 2014; Schummers et al., 2008; Sekiguchi et al., 2016; Thrane et al., 2012; Wang et al., 2006). In the ferret visual cortex, visually-evoked calcium responses in astrocyte somata is tuned to the orientation of a drifting grating, matching the receptive field properties of neighboring neural circuits (Schummers et al., 2008). In addition, localized calcium events are commonly observed to occur spontaneously throughout astrocyte processes (Agarwal et al., 2017; Nett et al., 2002; Rungta et al., 2016; Shigetomi et al., 2013), and their frequency can be modulated by the level of neuronal activity (Agarwal et al., 2017; Wu et al., 2014). These local events are likely to be of functional significance, as many of the consequences of astrocyte calcium remain intact in IP_3_R_2_ knockout mice, in which local, but not global, calcium signaling persists (Srinivasan et al., 2015). Even small portions of astrocyte branches likely contact synapses from hundreds of neurons, and while individual synapses have been shown to elicit small calcium events (Di Castro et al., 2011; Panatier et al., 2011), the interaction of multiple inputs have not been studied under physiological conditions. While astrocyte processes also respond to sensory stimulation *in vivo* (Otsu et al., 2015; Stobart et al., 2016; Wang et al., 2006), it is unknown whether different portions of an astrocyte are able to discriminate between different stimulus patterns which activate distinct pools of synapses. Ultimately, the answer to this question will define whether individual astrocytes should be considered as the relevant functional unit, or whether smaller functional subdomains exist within astrocytes and carry functionally relevant signals.

## Results

To address this question, we performed *in vivo* two-photon calcium imaging in young adult ferret visual cortical astrocytes while presenting orientated grating stimuli to the eyes (Fig 1A). Our virally-mediated strategy resulted in GCaMP6s expression throughout the entire cell of a sparse subset of cortical astrocytes (Fig 1B; Fig S1; Fig S4). Similar to previous reports (Shigetomi et al., 2013), AAV infection and GCaMP expression had no detectable effects on GFAP expression or cell morphology (Fig S1). We observed ongoing dynamic calcium activity in the absence of visual stimulation, comparable to that previously observed with similar approaches (see Video S1). Visual stimulation with drifting grating stimuli elicited reliable and robust responses (Fig 1C) that were similar to those obtained with organic calcium indicators (Schummers et al., 2008). Responses were time-locked to the stimulus onset, with a delay of 2.2 ± 0.15s (half rise time), peak time of 4.1 ± 0.11s, and duration of 3.9 ± 0.82s. Consistent with previous reports (Paukert et al., 2014; Schummers et al., 2008; Thrane et al., 2012), the amplitude and prevalence of calcium activity (both spontaneous and visually evoked) were strongly correlated with the global brain state measured by EEG (Fig S2). Specifically, calcium signaling was robust during desynchronized brain states, but nearly absent when large-amplitude, low-frequency EEG fluctuations were prominent.

**Figure 1.**
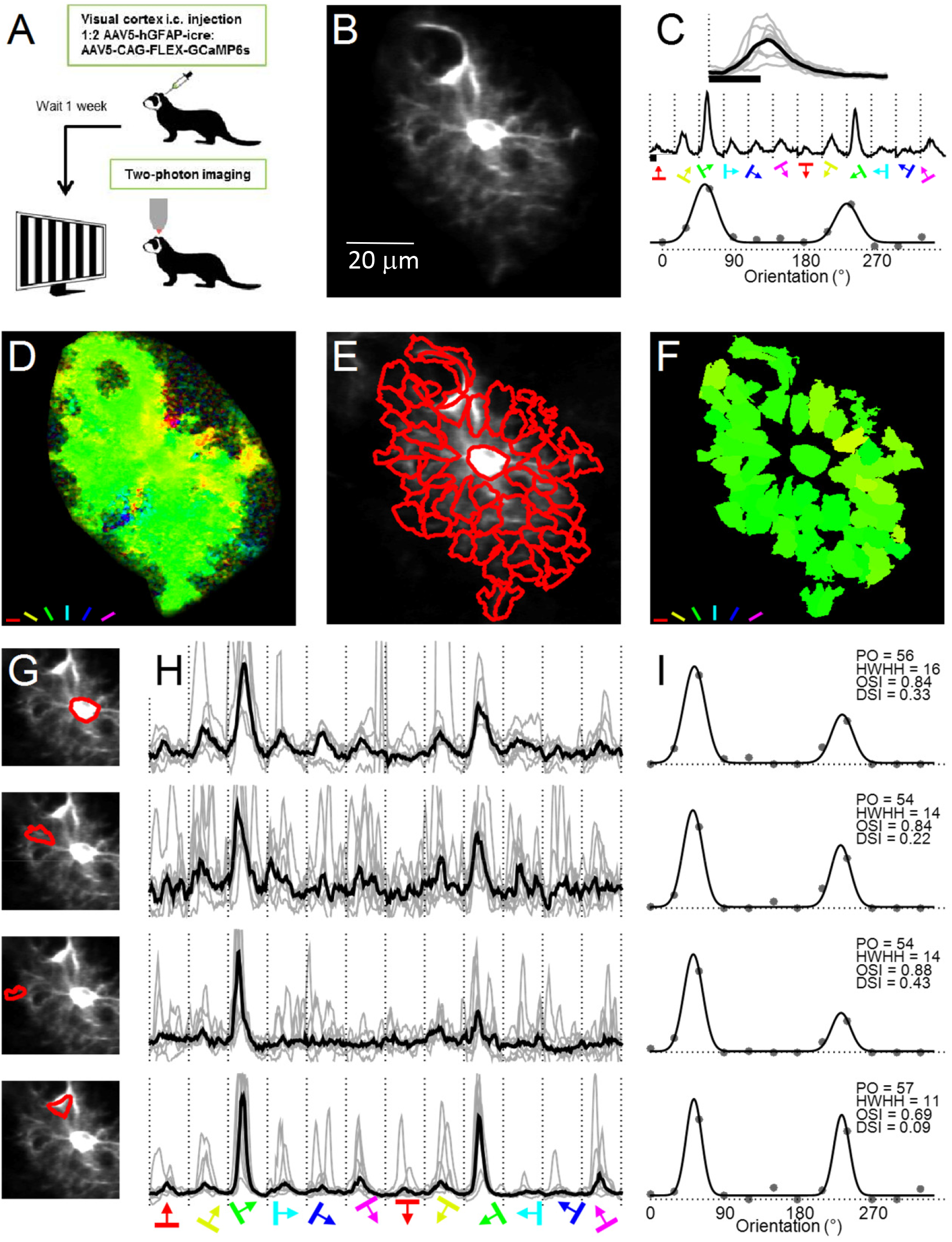
Orientation tuning of visually-evoked subcellular astrocyte calcium responses. **A.** Schematic of the experimental design. **B.** Structural image of an example astrocyte expressing GCaMP6s. **C.** *Top:* Individual trial traces (gray) and median over trials (black) for the astrocyte in **B** in response to presentation of preferred orientation visual stimulus (denoted by black bar). *Middle:* Trial averaged responses to each of 12 orientations. The onset of each stimulus is indicated by the dashed vertical lines. *Bottom:* Orientation tuning curve values computed from the evoked traces at each orientation (circles), and the fitted double-gaussian tuning curve (solid line). **D.** Pixel-based orientation preference map for the cell. **E.** Segments derived from ICA analysis, overlaid on the structural image. **F.** Segment-based map of preferred orientation. **G-I.** Quantitative analysis of orientation tuning in four segments (from top to bottom: soma, proximal process, distal process and endfoot). *G:* Segment outlines. *H:* dFF responses of each segment for 12 stimulus directions. *H:* Orientation tuning curves (gray dots) and fitted double-gaussian function (solid line) of each segment. PO: preferred orientation (in degrees); HWHH: half width of the gaussian at half maximal response (in degrees); OSI: orientation selectivity index; DSI: direction selectivity index.

As expected in ferret visual cortex (Schummers et al., 2008), the magnitude of the response depended strongly on the orientation and direction of motion of the grating stimulus (Fig1C). When response amplitude was plotted as a function of stimulus direction, the tuning curve was well fit by a double Gaussian function, which captures the preferred orientation (PO) at both directions of motion (Fig1C bottom). We observed strong responses with similar preferred orientation in most of the pixels throughout the territory of this astrocyte, which can conveniently be visualized in the pseudo-colored orientation pixel map (Fig 1D; Fig S3). The orientation preference map, obtained from a larger field of view containing this astrocyte, showed a gradual progression in preferred orientation, evident in both the traces of calcium activity (Fig S4C) and the orientation pixel map (Fig S4B). This spatial mapping of preferred orientation reflects the tuning of the underlying neural circuit, which is organized in a columnar orientation preference map in the ferret. Calcium responses of astrocytes separated by ~100 μm can respond to very different stimulus orientation (e.g. cells 1 and 3), indicating that they were triggered by activity in distinct populations of neurons.

These data suggest that the calcium responses in neighboring astrocytes are triggered independently by local neuronal activity. If each astrocyte responded to visual stimulation as a functional unit, we would expect individual astrocytes to have uniform preferred orientation throughout their territories, with distinct changes in PO at the boundaries between neighboring astrocytes. Alternatively, if local synaptic activity evoked local calcium responses within astrocyte branches, we would expect independent responses of different segments of individual astrocytes. To address this question, we analyzed the visually-evoked calcium responses in different segments within individual astrocytes. The boundaries of functional compartments of calcium signaling are not readily identified within the complex morphology of GCaMP6 filled astrocytes (Bindocci et al., 2017; Shigetomi et al., 2013). To overcome this, we identified functional segments within astrocytes by adapting the PCA/ICA segmentation approach (Mukamel et al., 2009), which defines segments of the image based on the spatial and temporal dynamics of the calcium activity itself, naturally dividing the astrocyte into its most independent segments, unbiased by experimenter input. This analysis generated some segments that correspond to distinct structural elements of the cell and others that encompassed regions of diffuse sub-resolution ‘gliopil’ with no visually resolvable morphological structure (Fig 1E). We then computed the time-course of calcium concentration for each of these segments for subsequent analysis of visual responses to the different stimulus orientations. Visual responses with significant orientation tuning were detected in 62 out of 64 segments. As expected from the pixel-maps (Fig 1D), the preferred orientation computed from these tuning curves was similar for all the segments (Fig 1F), however, the time-course of responses and other details of their responses, including the spontaneous events not time-locked to the stimulus and the degree of direction tuning varied from segment to segment (Fig 1G-I).

Neuronal responses in visual cortex display large stochastic variability of response amplitude from one trial to the next, which exhibits only modest correlation between neighboring neurons (Ecker et al., 2014; Goris et al., 2014; Lin et al., 2015). If the astrocyte calcium responses in localized segments of the cell are triggered independently by adjacent neural activity, we would predict that trial-to-trial differences in response amplitude would not be uniform throughout the cell, but would vary from one segment to the next. Indeed, we found that different segments of the same cell respond more strongly on different repetitions of the same (preferred) stimulus (Fig 2A,C), despite having similar orientation preference (Fig 2B). For example, segments 3 and 4 have a similar pattern of responses whereas segments 5 and 6 did not tend to have large responses on the same trials as each other (Fig 2C). Across all segments in the cell, a different complement of segments was active on each repetition of the stimulus (Fig 2D). The propensity to have highly variable responses did not have any discernable spatial organization with regard to location within the territory of the cell (Fig 2E). We quantified this pattern by computing the correlation across trials between each pair of segments in the cell (Fig 2F), which showed a broad range of correlation coefficients, albeit with a bias for positive correlations. Thus, not only are spontaneous, internally generated calcium events local and independent, but neuronally-driven ones also, which might be obscured when averaging across trials. When the same calculation was applied across all pairs of segments in all cells (n=25), the same trend was observed. From this correlation analysis alone, we were unable to distinguish whether the bias towards positive correlations reflected broad calcium events within the cell, or correlations of neural activity that triggered them. To address this issue, we analyzed the rare cases in which we were able to simultaneously measure responses of adjacent pairs of cells (Fig 2G,H). When we compared the correlation of segments within the same cell to those between neighboring cells, we found nearly indistinguishable distributions of correlation coefficients (Fig 2I). Together, these analyses demonstrate that visually-driven responses in individual segments do not simply reflect global events throughout the cell, but are largely independent of one another, regardless of which parent astrocyte they belong to.

**Figure 2.**
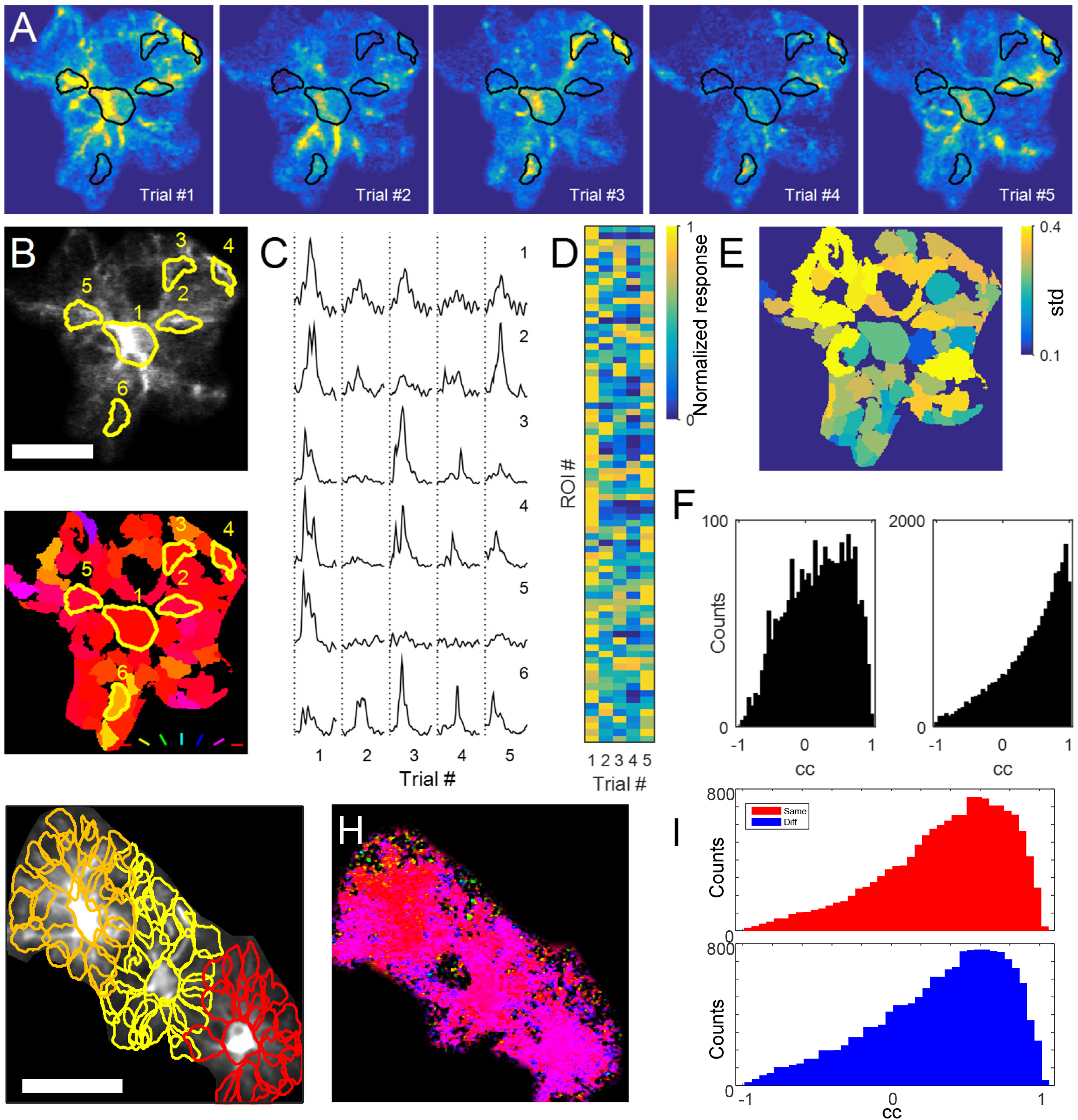
Variable subcellular spatial patterns of activity evoked by repeated stimulation. **A.** Response amplitude maps for an example astrocyte to five trials of the preferred orientation. Each image shows the maximal amplitude of the response at each pixel (dF; a.u.). Six example segments derived from ICA analysis (out of 79) are outlined in black. **B.** Overlay of example segments on the structural image (*top*). Orientation preference segment map (*bottom*). Scalebar is 25μm. **C.** Traces of the time course of response for the corresponding segments labelled in B to each of the five trials. **D.** Matrix of response amplitude for all 79 segments normalized to the maximal response for each segment across all trials. **E.** Spatial representation of the standard deviation of normalized response amplitude across trials for a each segment. **F.** Histogram of the correlation coefficient (cc) of response amplitude across trials between each of 2278 segment pairs for the same astrocyte (left). Population histogram of all 25,909 segment pairs for all 25 astrocytes (right). **G.** Structural image of three adjacent astrocytes, with ICA segments outlined for each cell. Scalebar is 50μm. **H.** Pixel based orientation preference map for astrocytes in **G. I.** Histograms of correlation coefficients computed between pairs of segments within the same astrocyte (red), or pairs in different cells (blue).

The orientation preference map contains regions where orientation preference changes gradually and slowly, as well as regions, such as pinwheel centers, where orientation preference changes rapidly on the scale of tens of microns. If astrocyte segments respond independently to nearby synapses, then in rapidly changing regions synaptic activity impinging on different parts of an astrocyte would have different orientation tuning, which we predicted would be reflected in the astrocyte calcium signals. We tested this hypothesis by searching for astrocytes located in high change regions of the map. As expected (Schummers et al., 2008), we found that in these regions of the map, adjacent astrocytes had different preferred orientations (Fig 3A), demonstrating that different astrocytes respond to the activity of different neurons. To test whether different segments within an individual astrocyte responded to different orientations, we performed high-resolution scans of an astrocyte located within this transition zone. When we examined the orientation tuning within a single astrocyte, we found that indeed different segments of the same astrocyte had different preferred orientations (Fig 3B-C). On the flanks of the tuning curves, orientations that elicited strong responses in one segment elicited barely detectable responses in other segments, despite being separated by only tens of microns within the same cell (Fig 3C). Furthermore, the overall pattern of preferred orientations within this cell corresponds to the pattern of the orientation preference map surrounding it. This suggests that not only can different branches respond independently, but that they respond to distinct information content contained only tens of microns apart in the neural circuit.

**Figure 3.**
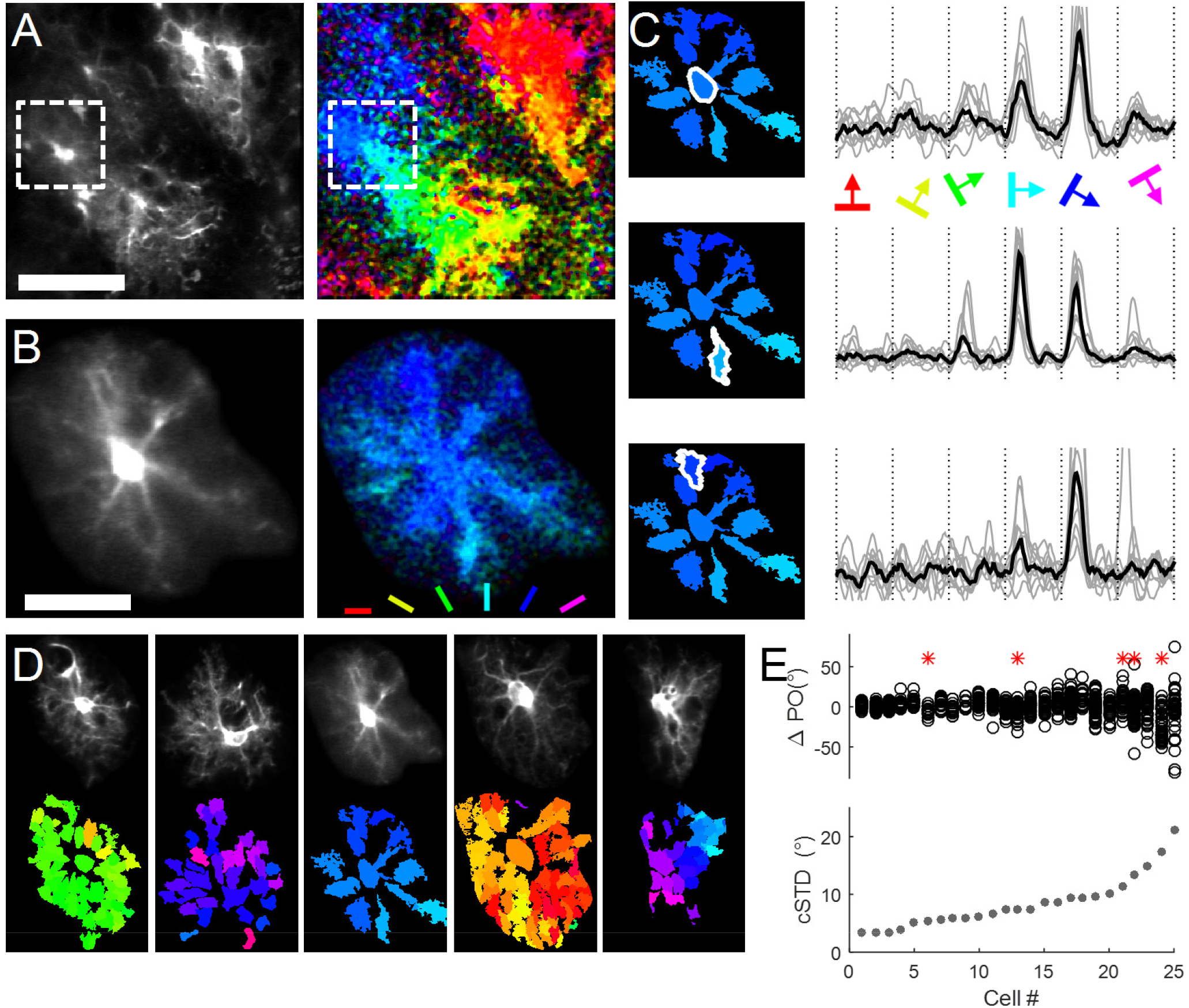
Different orientation tuning of segments within individual astrocytes. **A.** Structural image and orientation preference map from a field of astrocytes. Scalebar is 100μm. **B.** Morphological image and orientation preference pixel map for one astrocyte located in a high rate-of-change region of this FOV, indicated by the box in **A**(~50 microns deeper). Scalebar is 25μm **C.** Segment based orientation preference map and traces from three example segments within the cell. This cell was highly direction selective, so only 180 degrees are shown. Different preferred orientations of segments within the cell are apparent. **D.** Segment-based maps for five astrocytes demonstrating the range of preferred orientation uniformity encountered within individual cells. The cells are sorted from more uniform preferred orientation on the left, to more diverse orientation preferences on the right. **E.** Quantification of the spread of POs within each cell. *Top:* Scatter plot of the preferred orientation for each segment within each cell. The PO is normalized relative to the PO of the soma for visualization. The cells are sorted from least to most spread of PO (cSTD). The example cells in **D** are indicated by *. *Bottom:* The circular STD for each cell is shown by the circles, in ascending order.

Across our population of astrocytes, we observed cells with uniform preferred orientation across all segments, as well as cells with systematically changing preferred orientation tuning (Fig 3D). We quantified the uniformity of preferred orientation of each astrocyte by first computing the difference in PO for each segment, relative to the soma (Fig 3E, top), and then computing the circular standard deviation (cSTD) of the POs of all of its segments, for each cell (Fig 3E, bottom). This analysis demonstrated that while many cells had uniform PO, there was a broad distribution of cSTD, including several cells with large dispersion of PO among their segments. Other quantitative measures of tuning curves, including direction selectivity and tuning bandwidth – which would impact information coding - also showed variable diversity across different segments across the population of cells (Fig S5).

We next asked whether this skewed distribution of cSTD in astrocyte segments reflects the distribution of local dispersion of neuronal responses that arises from the periodic pinwheel structure of the orientation preference map. In order to evaluate this interpretation, we generated a model of the spatial interaction between astrocytes and the orientation preference map. We obtained high-resolution, large field of view, maps of orientation preference at sub-cellular scale, using two-photon imaging of neurons expressing GCaMP6s (Fig 4C), which includes signal from dendrites, axons and cell bodies. We then generated a model astrocyte, with geometry, segment number and size based on our data (Fig 4A; Fig S6). We simulated the PO for each segment in the model astrocyte by taking the average PO value of all the pixels in the neuronal orientation map that the segment overlaps (Fig 4B), and computed the cSTD for the model astrocyte. Consistent with our astrocyte imaging data, the model produced a wide range of cSTD values, with locations in smooth portions of the map generating low cSTD values (Fig 4B left), and locations near pinwheel centers generating high cSTD values (Fig 4B right). When we quantitatively analyzed the distribution of the cSTD values from 100 repetitions of 25 randomly placed simulated positions, we found a strong correspondence between the model and the data (Fig 4D), suggesting that the distribution of cSTD values in the data reflect the structure of the neural circuit. The results of the model therefore support the interpretation that independent subdomains within astrocytes integrate local neuronal activity independently.

**Figure 4.**
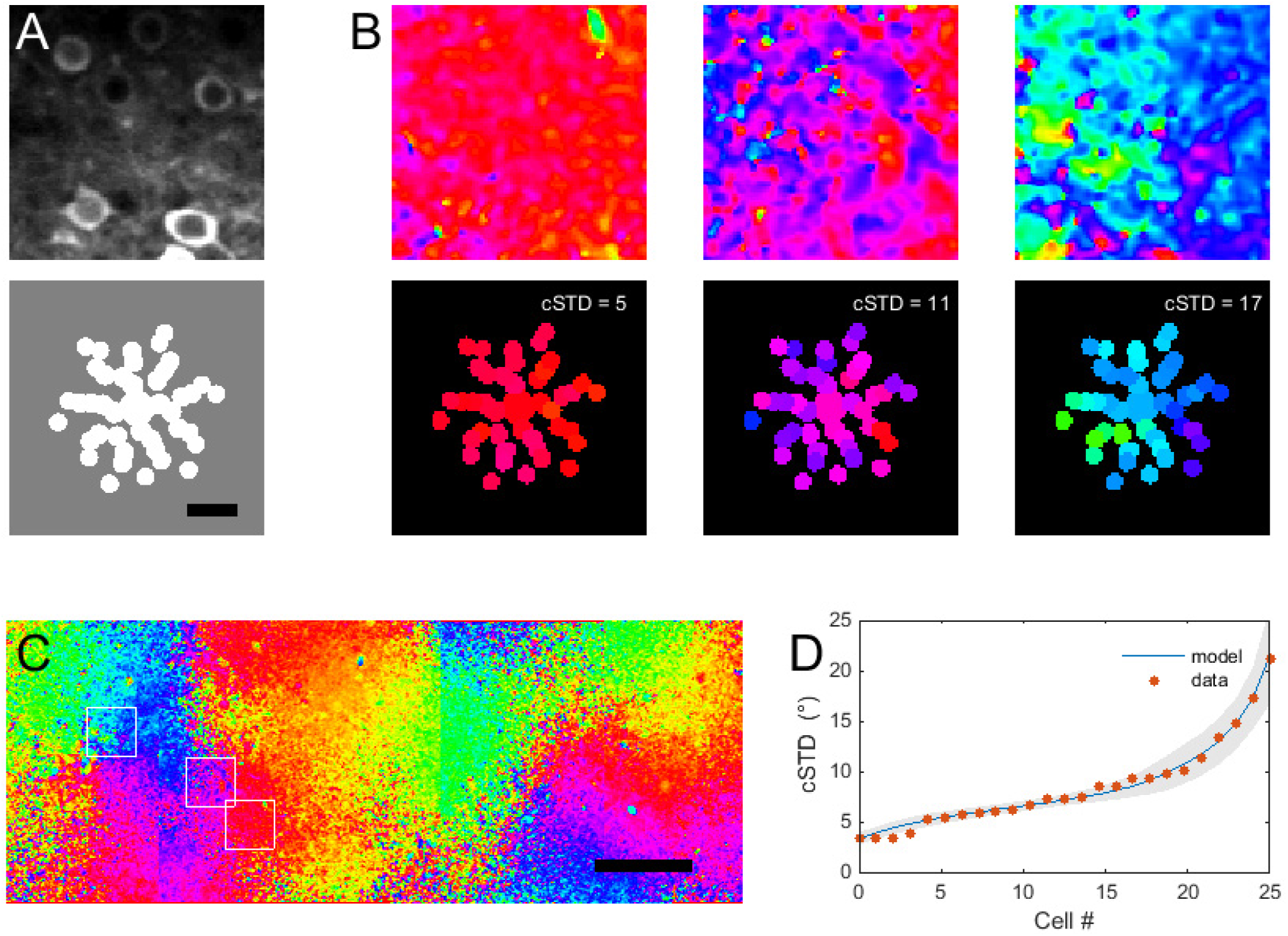
Linear astrocyte integration model captures statistics of astrocyte diversity. **A.** Depiction of model astrocyte (*bottom*) and an example FOV of GCaMP6s-expression in neurons (*top*; same scale). **B.** Three representative segment maps (*bottom*) from the model astrocyte at different locations in the orientation map (*top* - regions marked in **C**). **C.** Montage of tiled orientation preference pixel map generated from neuronal GCaMP6s imaging. Pixels with weak responses or low SNR were removed. **D.** The average (+/- sem) sorted distribution of cSTD for all simulated astrocyte locations (n = 100 repetitions of 25 random locations) overlaid on the cSTD values from all imaged astrocytes (n = 25).

## Discussion

We have measured the local calcium responses of distinct segments within the territories of cortical astrocytes *in vivo* during visual stimulation. Making use of the columnar organization of orientation preference in the ferret visual cortex, we have demonstrated that different segments of the same astrocyte respond independently to different visual stimuli, forming functional subdomains. The spatial scale of these functional subdomains was 5-10 μm, indicating that astrocyte calcium signals carry information about visual stimulus identity with high spatial precision. As a result, for astrocytes located near transition zones, where the representation of orientation changes rapidly among nearby neurons, different subdomains within a single astrocyte represent distinct information about the visual stimulus.

Our results suggest that a major component of astrocyte calcium signaling is triggered by very local synaptic activity, under naturalistic activation of neural circuit activity *in vivo*. While a direct demonstration that local synaptic activity triggers the events we observe is not technically feasible *in vivo*, this is the most plausible interpretation, based on the known circuitry of ferret visual cortex. Both glutamatergic and GABAergic transmission are orientation tuned and observe the requisite spatial organization to account for the spatial specificity of orientation tuning we observe (O’Herron et al., 2016; Wilson et al., 2017), and both could plausibly drive astrocyte calcium responses (Mariotti et al, 2018). Neuromodulatory systems, which may contribute to some portion of the spontaneous local or global calcium activity we observed (Chen et al., 2012; Ding et al., 2013; Paukert et al., 2014; Takata et al., 2011), have not been shown to have the response characteristics or spatial precision to drive the orientation tuned visual responses in astrocyte subdomains, consistent with a more spatially diffuse gating (Papouin et al., 2017; Paukert et al., 2014) of the local events that we describe.

It is likely that even the smallest visually-evoked calcium responses that we observe result from pooling of synaptic activity from multiple synapses. This conclusion is supported by several lines of evidence. First, the ER is positioned throughout a large portion of the processes, but not into the finest processes, at which individual synapses are contacted (Patrushev et al., 2013). Though the mechanism(s) of synaptic signaling that ultimately trigger release from the ER remain to be elucidated (Shigetomi et al., 2016; Volterra et al., 2014), this arrangement would likely require cooperative activation of multiple synapses to trigger release from internal stores. Second, the size of the smallest visually-driven domains that we observed is substantially larger (~5 μm) than would be expected from single-synapse activation, based on *in vitro* brain slice studies (Di Castro et al., 2011; Panatier et al., 2011). This is not due to insufficient ability to resolve smaller events, as we routinely observed micron-scale spontaneous events that were not triggered by visual stimulation (See Supp. Movie 1). Whether these micro-domains reflect spontaneous vesicle fusion, spontaneous activation of IP3 hot-spots, mitochondrial events, or some other cell-intrinsic or synaptic process remains an area of ongoing investigation (Rusakov, 2015). Furthermore, the amount of scatter in the synaptic organization of orientation preference in the ferret (Wilson et al., 2016), would predict more scatter than we observe if the astrocyte calcium signals we detected reflect monitoring of individual synapses. Our model succeeded in recapitulating the statistics of astrocyte orientation tuning despite using a relatively simple rule for predicting the PO of astrocyte segments: linearly averaging the PO of the surrounding neural activity. That such a simple model can predict the astrocyte calcium responses suggests that astrocyte subdomains may act as spatial low-pass linear filters of neuronal activity.

The factors that determine the boundaries of functional subdomains, which might include spatial restrictions on biochemical diffusion as in neurons (Colgan and Yasuda, 2014; Yuste et al., 2000), non-uniform clustering of receptors (Buscemi et al., 2017) or calcium stores, such as ER (Patrushev et al., 2013) or mitochondria (Agarwal et al., 2017), remain to be fully explored. It may be interesting for future studies to evaluate whether the boundaries are fixed, across different stimulus conditions (e.g. visual stimulation vs neuromodulatory input), during development or plasticity. An interesting possibility is that astrocytes function at multiple spatial scales in order to modulate different aspects of neuronal activity. For example, it is possible that the capacity to modulate individual synapses may be limited to low activity regimes (Di Castro et al., 2011; Panatier et al., 2011), where synaptic activity is sparse, and the capacity to synchronize populations of neurons (Fellin et al., 2004; Pirttimaki et al., 2017) may become prevalent under conditions of high activity which engage cell-wide calcium events in astrocytes. Our data indicate that under physiological *in vivo* conditions, in which the circuit is engaged by a natural sensory stimulus, intermediate scale functional subdomain activity predominates. Recent demonstrations that parallel gliotransmission pathways can exist within single astrocytes (Schwarz et al., 2017) lend support to the notion that multiple signaling pathways are available to enable flexible interactions with neurons under different conditions.

In the visual cortex of the ferret, in which visual information content is precisely arranged spatially, different subdomains take part in processing different functional information. It is likely that this principle applies to other cortical circuits with columnar mapping of sensory receptive field properties, and neural circuit organization may help to explain why astrocyte calcium responses are more reliably reported in mouse somatosensory than visual cortex (Lind et al., 2013; Takata et al., 2011; Thrane et al., 2012; Wang et al., 2006; though see Sonoda et al., 2018; Slezak et al., biorxiv). In other brain regions or species, where functional information is not as strongly spatially organized, the functional significance of local astrocyte signals will require more investigation. It is also worth noting that our scanning technique only allows a two-dimensional view of an astrocyte, and we are undoubtedly missing events out of the focal plane (Bindocci et al., 2017). However, because the orientation preference map is also two-dimensional - in the same plane as our imaging plane – our data would be expected to capture the relevant relationship between neuronal and astrocyte activity.

While the functional significance of these signals for neural circuit function, vascular regulation, ionic homeostasis or energetic supply remains to be elucidated, our results suggest the possibility that local signaling is a predominant feature of astrocyte calcium signaling. The recent demonstration of local protein translation within distal astrocyte processes (Sakers et al., 2017) broadens the range of regulatory mechanisms that could be triggered by local calcium events. Future technical advances that enable simultaneous subcellular manipulation of astrocyte activity and monitoring of the resulting impact on neuronal activity will be required to understand the importance of these signals for information processing.

## Materials and Methods

### Animals

Experiments were performed in young adult (P70-P150) female ferrets (*Mustela putorius furo;* Marshall Farms). All experiments were performed in accordance with NIH guidelines for the Care and Use of Laboratory Animals and approved by the Max Planck Florida Institute for Neuroscience IACUC.

### Surgery and virus injection

Ferrets were given atropine (0.06mg/kg) and slow-release buprenorphine (0.6mg/kg) peri-operatively and anesthetized with ketamine (25 mg/kg)/xylazine (1.5 mg/kg). The eyes were protected with ophthalmic ointment during the surgery and moistened afterward with silicone oil. Hair was clipped, and EKG, etCO2 and spO2 were monitored thorough out the experiment to monitor the depth of anesthesia. Core body temperature was maintained at 38.3 degrees using a feedback-regulated heating blanket and an IV line was placed in the cephalic or saphenous vein for fluid delivery. The animals were intubated, and mechanically ventilated with isoflurane 1.5-2.0% in a 1:1 mixture of O_2_ and N_2_O. Ferrets were mounted in a stereotaxic frame, the surgical site was prepared under aseptic technique and bupivacaine was applied subcutaneously. An incision was made and skin, fascia and muscle overlying the left visual cortex were retracted to place a titanium head plate to the skull with dental acrylic. A craniotomy (~5mm) was made using a high-speed burr drill and the dura mater was removed to expose the brain. In some cases, two skull screws were implanted over the frontal lobe to enable EEG recording. Virus was mixed with fast green to visualize the spread of the injection through the surgical microscope. The virus was injected (250-500nL in 3 to five injection sites) into visual cortex at a depth of ~250 microns (layer 2/3) in increments of 18nL every minute using a beveled fine glass pipette (20-40 micron tip diameter) attached to a NanoJect. The pipette was retracted 10 minutes after the end of each injection. After virus injections, the craniotomy was filled with agarose (2%) dissolved in ACSF, and the window was closed with a glass coverslip and cemented (Metabond) to the skull to seal the craniotomy. The ferrets were then allowed to recover from anesthesia and returned to their home cage. Two-photon imaging was performed after period of 6-14 days.

### Viral transduction

We used a cre-recombinase system to drive GCaMP6s expression in astrocytes. For this, AAV5-GFAP2.2-cre (Vector Labs) and AAV5-CAG-FLEX-GCaMP6s (Penn Vector Core) were injected together (1:2). This strategy enabled sufficient expression and adequate brightness of the calcium indicator for in vivo imaging. At the times used in this study (6-14 days), GCaMP expression was nearly exclusively observed in astrocytes (confirmed by post-hoc immunostaining in a subset of ferrets) and occasional neuronal expression was observed. Neurons were readily discriminated by their morphology and time-course of changes in calcium signal and avoided when choosing a field of view for imaging. At longer expression times, neuronal expression became more prominent, which limited the time points at which we could image to less than 15 days post-injection. To image neurons, GCaMP6s expression was driven with either AAV1-hsyn-GCaMP6s or AAV9-hsyn-GCaMP6s (Penn Vector Core).

### Calcium imaging

On the imaging day, ferrets were anesthetized with isoflurane (1.5-2% in a 1:1 mixture of O_2_ and N_2_O) intubated and mechanically ventilated. An IV line was placed in the cephalic or saphenous for fluid delivery and EKG, etCO2, spO2, and rectal temperature were continuously monitored to ensure the animal was within normal physiological condition, and the depth of anesthesia was adequate. The ferret was then mounted into a frame using the head-mounted headplate, and placed onto the stage of the two-photon microscope. Eyes were treated with atropine and phenylephrine drops and the left (ipsilateral) eye was covered to ensure monocular stimulation. To prevent spurious drifts in eye position, they were paralyzed with pancuronium bromide 0.2 mg/kg/hr. In ferrets with implanted skull screws, EEG leads were connected.

Two-photon imaging of GCaMP6s was performed with an Ultima IV microscope (Prairie Technologies, Middleton WI), coupled to a titanium/sapphire laser providing excitation at 1000nm with ~100 fs pulses at 80 MHz (Tsunami; Spectra-Physics, Menlo Park, CA). The emitted fluorescence was detected using GaAsP photomultiplier tubes (Hamamatsu, Shizouka, Japan) through a 16x, 0.8 NA objective lens (Nikon). The PMT gain and laser intensity were set to achieve adequate signal-to-noise ratio while avoiding excessive heating or phototoxic effects or artifacts, typically ~600V and ~20 mW, respectively. For high magnification scans of individual astrocytes, time series images were collected at 4 Hz (2 Hz for 2/25 cells). Optical zoom was set between 6.6 and 12, yielding a FOV of 72-123 μm, with corresponding pixel sizes of 0.2-0.48. Larger FOV (215-430 μm) scans containing multiple astrocytes were obtained with optical zoom or 2-4x zoom were acquired at 2Hz. Neuronal orientation maps were acquired with similar parameters, and multiple tiled FOVs were stitched together in Matlab for analysis.

### Visual stimulation

A computer monitor was placed ~30 cm in front of the ferret eyes for visual stimulus presentation. The monitor and ferret were draped to prevent any light from the stimulus monitor from reaching the photodetectors. Visual stimuli were generated by custom routines written in Matlab, making use of the Psychophysics Toolbox (Brainard, 1997). Visual stimuli consisted of high contrast (100%) square wave gratings (spatial frequency 0.06-0.12 cpd; temporal frequency 2.0-5.0 cps) which orientation ranges between 0-360 degrees in increments of 22.5 or 30 degrees. Each stimulus was presented for 4 seconds, with an inter-stimulus interval ranging from 4-28 seconds, during which an equiluminant gray screen was presented. Visual stimulus and data acquisition were synched via a TTL pulse, with temporal resolution of less than one monitor refresh (~7msec at 144Hz).

### Data analysis

Image files (tiff) acquired in PrairieView software were loaded into Matlab for all subsequent analysis. Frames were rigidly aligned to one another in the frequency domain (Ringach et al., 2016) to reduce the effects of respiration-induced motion. Image sequences were smoothed with a three pixel sliding boxcar filter in both spatial and temporal dimensions, and with a three pixel spatial median filter to remove spurious pixels, and manually cropped around the edges of the astrocyte territory to minimize spurious analysis of system noise outside of the astrocyte. Only astrocytes that were visually responsive for at least three trials (average of six) were accepted for further analysis.

To compute pixel maps, all trials were averaged, and the response amplitude of every pixel was computed for each orientation to generate response amplitude maps (see Fig 2A and Fig S3). The amplitude was calculated as the maximum value during a user-defined time window encompassing the response time-course minus the median of the values during a user-defined time window after the response had returned to baseline. These pixel values were smoothed in space and orientation with a boxcar convolution to remove pixel noise. The preferred orientation was computed as the vector average, and the response amplitude was computed as the maximal response across orientations, in units of dF/F_0_ (see Fig S3).

Astrocyte segmentation was done using the PCA/ICA approach developed by Mukamel et al., (2009) (Mukamel et al., 2009). Temporal and spatial components were given equal weights and the number of ICAs was chosen as the intersection of the PCA curve with 2 times the noise. Only the spatial components of the recovered components were used to define the spatial astrocyte segments. The occasional small spatial overlap of multiple segments was removed, and any segments smaller than 50 pixels were discarded. Due to variation in the exact position of the soma relative to the large proximal processes at the chosen z-plane, the soma segment was manually defined, and any ICA segments that overlapped the soma were discarded. Defined in this way, astrocytes (n = 25) had 50 ± 5 segments, with an average size of 7.2 ± 2.1 μm (equivalent circular diameter; n = 25 astrocytes from 12 ferrets), of which, 42 ± 4 (85%) had significantly orientation-selective visual responses, defined as described below.

The time series for each segment was then computed from the average value of all pixels within the segment, and was converted into units of dF/F_0_, using the 25^th^ percentile value across all time points as the F_0_ value. Unresponsive trials, presumably due to changes in brain state (see Fig S2), were removed based on visual inspection. The response amplitude for each stimulus was defined as the maximum of the response during a user-defined on period minus the median of the values during the off time when responses returned to baseline. Tuning curves were computed from these response amplitudes for each stimulus orientation, and least-squares fits of circular double-Gaussian functions were used to compute preferred orientation. Significant tuning was defined as a correlation coefficient of >= 0.75 between the data and the fits, and segments without significant tuning (15%) were discarded from further analysis. Response onset and duration were calculated from the half-rise time and half-decay times of 10X interpolated response profiles to the preferred orientation. OSI of tuning curves was defined as the magnitude of the vector average, and ranges from 0 (no orientation selectivity) to 1 (highly selective). DSI was defined as the difference between the response to the preferred and non-preferred directions divided by their sum, and ranges between 0 (equal response to both directions) and 1 (response to only one direction).

### Geometric astrocyte model

A model astrocyte was generated from the statistics of the segments obtained by the PCA/ICA segmentation. The geometry was a simplified version of the typical geometry of astrocytes based on the average sizes and distances of segments from the soma. The resulting segments in the model astrocyte were 41 circles in concentric rings around the soma with the following parameters: Distances: [9, 13, 18, 22, 27, 31, 36 μm]; Radii: [4.2, 4.0, 3.9, 3.8, 3.6, 3.6, 3.6 μm]; Number: [4, 6, 9, 9, 7, 6]. We performed a simulation to mimic the experimental data acquisition, by placing 25 astrocytes at random locations throughout the neuronal orientation preference map. This simulation was repeated 100 times. The preferred orientation for each segment was computed as the mean preferred orientation of all the pixels in the orientation preference map with which it overlapped. As with the analysis of the imaging data, the circular STD was then computed for each position, to model an astrocyte located at that position in the map.

### EEG recording and analysis

In a subset of experiments, EEG data was recorded simultaneously with calcium imaging via differential recording between two skull screws placed over the frontal lobe. Signals were amplified 2000X, bandpass filtered between 0.1-300Hz (Grass P511 amplifier), digitized at 1kHz and synchronized to calcium imaging data with 10s resolution. Power spectral analysis was used to designate each trial as synchronized or desynchronized, based on the ratio of power in low frequency (<10Hz)/high frequency (>10Hz) bands.

### Histology and Immunostaining

Ferrets were deeply anesthetized with sodium pentobarbital (Euthasol) and perfused transcardially with 50 ml of heparinized saline followed by 400 ml of fixative solution (4% paraformaldehyde in 0.1M sodium phosphate buffer saline (PBS), pH 7.4). Brains were removed and keep it in fixative solution at 4° C for 24 hours. 50μm parasagittal sections were obtained in 0.1M PBS (pH 7.4) using a vibrating microtome and stored at 4°C. Brain sections for immunoprocessing were incubated 60 minutes at room temperature in 5% normal goat serum (NGS, Jackson ImmunoResearch, cat# 005-000-121) to block non-specific labeling and incubated 48 hours at room temperature in a cocktail of primary antibodies and diluent (1% NGS in PBS containing 0.25% Triton X-100, pH 7.4). Primary antibodies were as follows: rabbit anti-S100β (Abcam, cat# ab868,1:500), goat anti-GFP (cat# ab5450, 1:2000) and chicken anti-GFAP (cat# ab4674, 1:500). After the primary incubation, sections were washed in PBS (5 × 3 minutes) and then incubated 3 hours in a cocktail of secondary antibodies conjugated to Alexa Flour dyes (Life Technologies) to tag the primary antibodies at a concentration of 1:500. Goat anti-rabbit Alexa 488 and Goat anti-mouse Alexa 594 were used to detect S100β and NeuN. Following secondary antibody immunostaining, tissue was washed and then incubated in Hoechst solution (1:20,000) and saline (pH of 7.4) for six minutes and washed 3 times in PBS to stop Hoechst staining. Stained sections were mounted onto Superfrost (VWR, cat# 48311-703) slides, coverslipped using Slow Fade Gold Antifade media (Life Technologies, cat# S36936) and stored at 4 ° C.

### Confocal Imaging

Fixed tissue was imaged with a Zeiss LSM780 confocal microscope (Carl Zeiss MicroImaging) with a Zeiss plan-Apochromat 40x/1.3 Oil objective (NA 1.3). Fluorophores were excited with an Argon laser 458, 488, 514 nm (for 488 nm), HeNe laser 594 nm (for 594 nm) and laser Diode 405 nm (for 405 nm).

## Acknowledgements

This work was funded by the Max Planck Florida Institute for Neuroscience and NEI R01EY02697 (J.S.). For the use of GCaMP6s, we gratefully acknowledge V. Jayaraman, R.A. Kerr, D.S. Kim, L.L. Looger and K. Svoboda from the GENIE Project, Janelia Farm Research Campus, Howard Hughes Medical Institute. We are grateful to Walter Hoover for performing immunohistology, and for surgical assistance.

## Data availability

The datasets generated during the current study are available from the corresponding author on reasonable request.

## Code availability

All custom Matlab code used in the analysis of the data and production of the figures will be made available upon reasonable request to the corresponding author.

---

**Figure S1.**
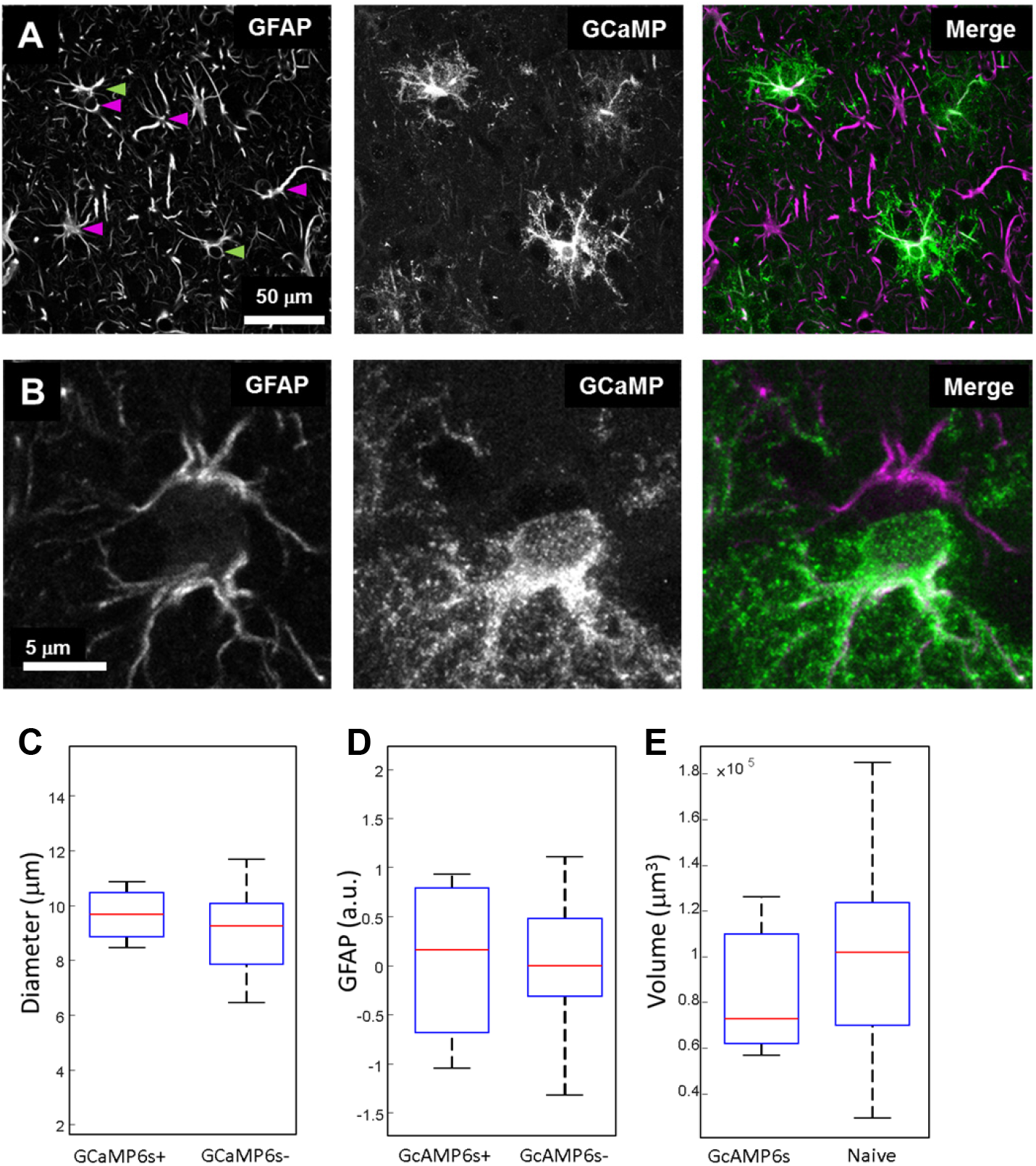
Lack of reactive gliosis in AAV-infected, GCaMP6s-expressing astrocytes. **A.** Two channel confocal image of fluorescent immunostaining for GFAP and GCaMP6s (GFP). GCaMP6s is only detectable in a sub-set of GFAP+ astrocytes. GFAP expression levels and pattern are similar in GCaMP6s+ and GCaMP6s− cells, some of which are indicated by green and magenta arrows, respectively **B.** Higher magnification images of two adjacent (‘kissing’) astrocytes. Visually similar GFAP expression is seen despite GCaMP6s expression being limited to only one cell. **C.** Quantitation of the cell body diameter of GCaMP6+ and GCaMP6-astrocytes assessed from double stained tissue as in A-B. Median values: 9.7 vs 9.3μm; p = 0.44, Wilcoxon test. **D.** Quantification of GFAP staining intensity of GCaMP6+ and GCaMP6-astrocytes. Median 0.16 vs 0.00a.u.; p = 0.9347, Wilcoxon test **E.** Quantitation comparison of the territory volume computed from z-stacks acquired *in vivo* for GCaMP6+ cells vs from cells electroporated with dextran-conjugated Alexa dyes in naïve ferrets (data from Lopez-Hidalgo et al., 2017) Median = 101,679 vs 72,700 μm^3^; p = 0.38, Wilcoxon test. For boxplots in C-E the central mark is the median, the edges of the box are the 25^th^ and 75^th^ percentiles, the whiskers indicate the data range.

**Figure S2.**
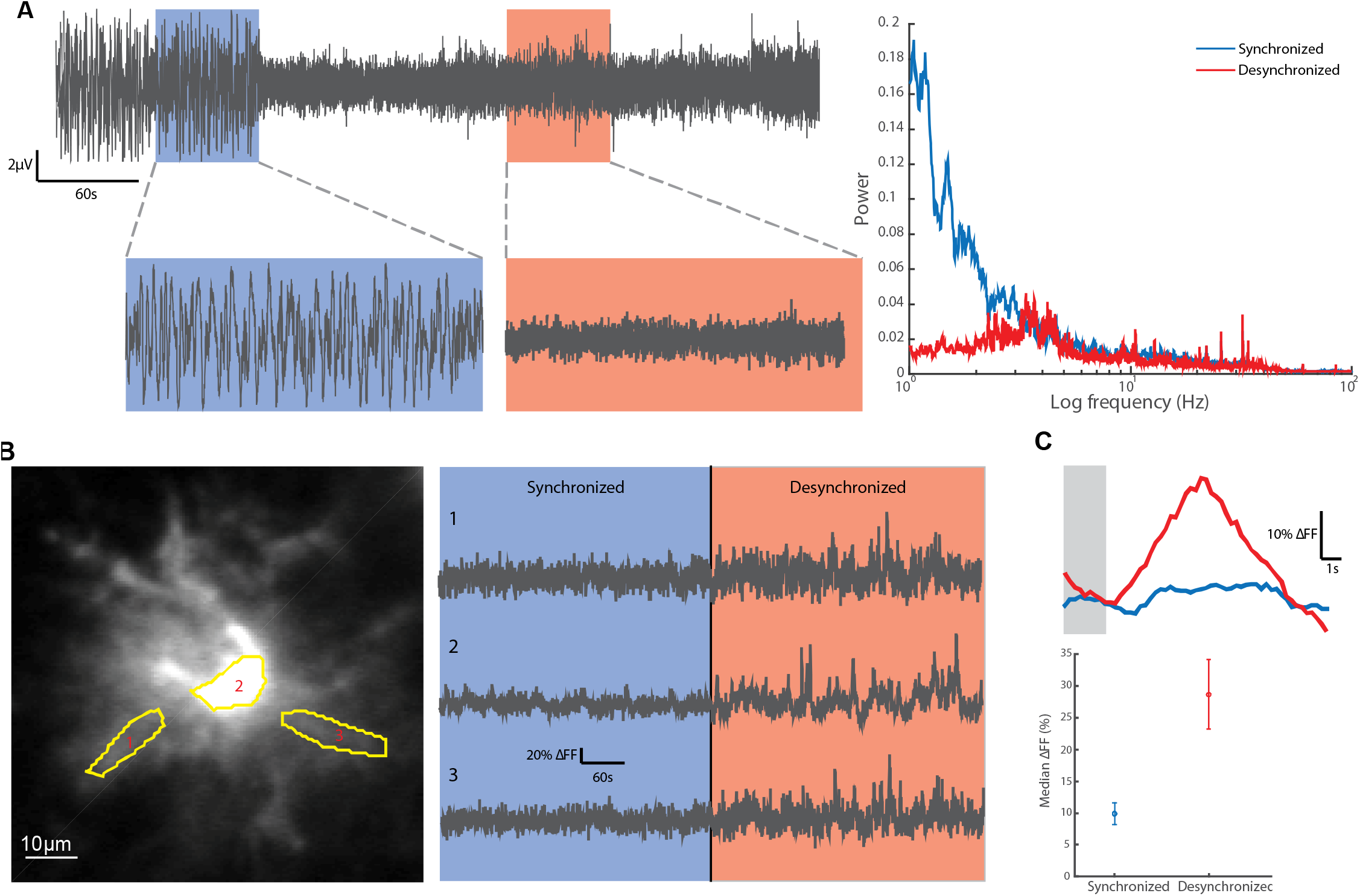
Brain-state dependence of astrocyte calcium signaling. **A.** *Left, e*xample EEG trace that demonstrates the transition between a state dominated by large amplitude, low frequency waveforms, which we refer to as the synchronized state (blue), and small amplitude, high frequency waveform, indicative of a desynchronized state (orange). *Right,* power spectra from the two states demonstrates distinct characteristic features. **B.** Astrocyte calcium activity from an example astrocyte during synchronized and desynchronized brain states. The traces from three segments, indicated in the left panel are shown in the right panel. These traces include epochs with the stimulus on, as well as with the stimulus off, and thus show both spontaneous and visually-evoked calcium activity. **C.** Analysis of the amplitude of visually-evoked calcium responses. *Top,* median over trials of the calcium response in all ROIs within the astrocyte in **B** in response to a 4s visual stimulus in the preferred orientation (gray bar) during the synchronized (blue) and desynchronized (red) phases. Bottom, median ± MAD of maximal amplitude to preferred orientation stimulus over ROIs in the cell presented in B.

**Figure S3.**
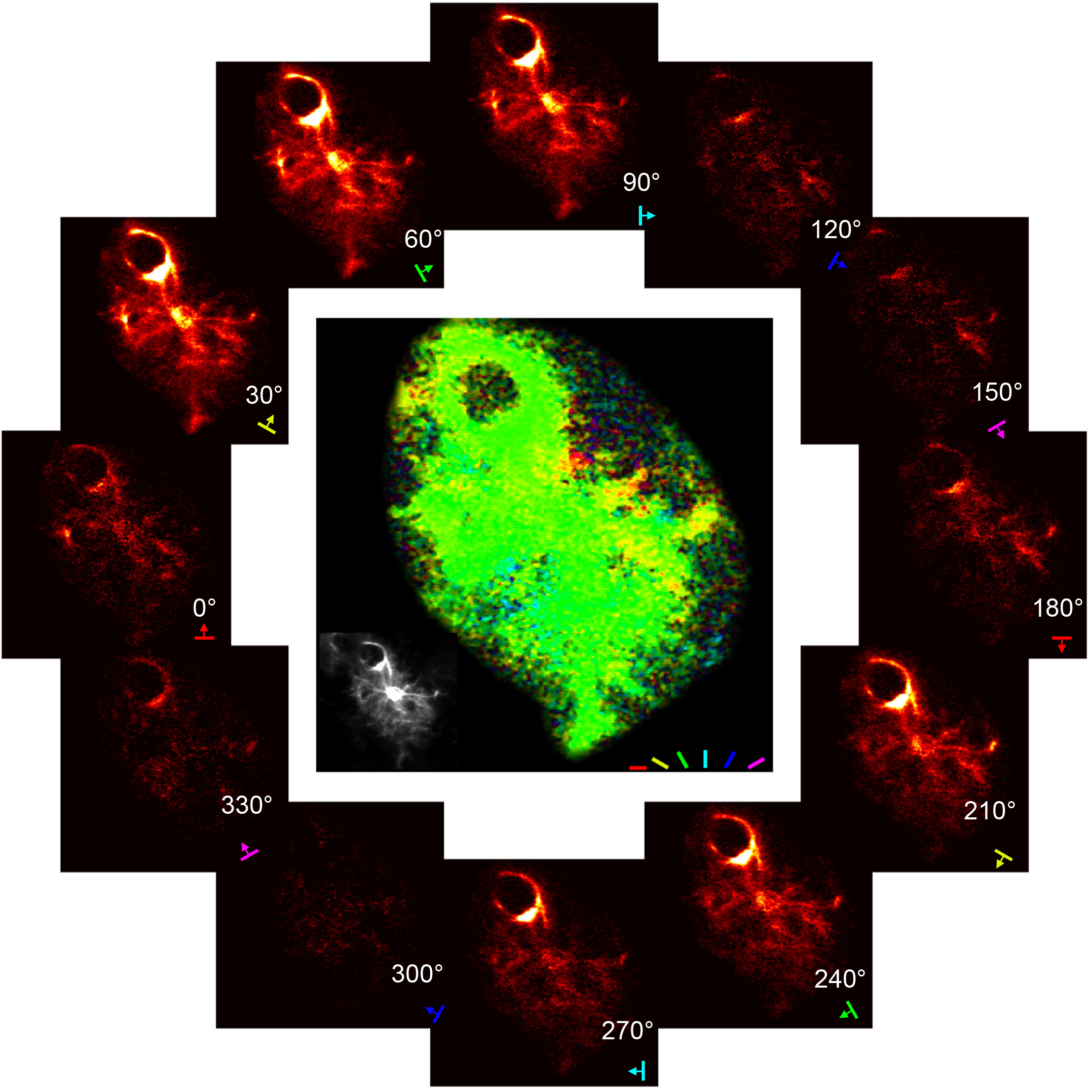
Computation of the pixel-based orientation preference map. The image for each orientation shows the change in fluorescence for each pixel during response to the indicated orientation (dF). Central image shows the pseudo-color coded pixel-based orientation preference map, where the color of the pixel indicates the preferred orientation and the color saturation depicts the strength of the response. The mean intensity image, demonstrating the morphology of the cell is shown in the inset. Same cell as in Fig 1.

**Figure S4.**
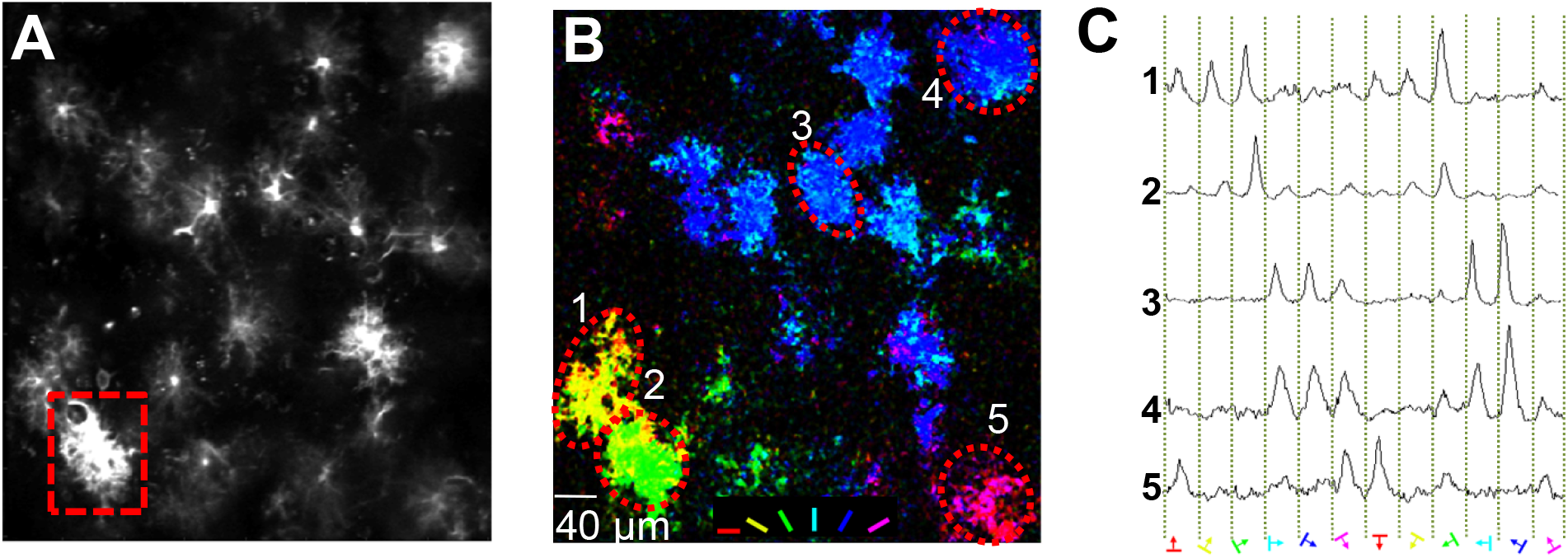
Orientation preference maps of astrocyte tuning. **A.** Structural image of an example astrocyte morphology obtained by two-photon imaging in vivo. Dashed box indicates the cell shown in Figure 1. **B.** Pixel-based orientation preference map for the FOV. The color of each pixel represents the preferred orientation computed from the responses of that pixel to all 12 orientations, according to the pseudo-color code. The brightness of each pixel indicates the amplitude of the response. **C.** Trial averaged responses to each of 12 orientations spanning 360 degrees demonstrating orientation and direction tuning for the five cells indicated in **B.** The times when stimuli were turned on are indicated by the dashed vertical lines. The orientation and direction of motion of each stimulus is depicted below, and also indicated by the pseudo-color code.

**Figure S5.**
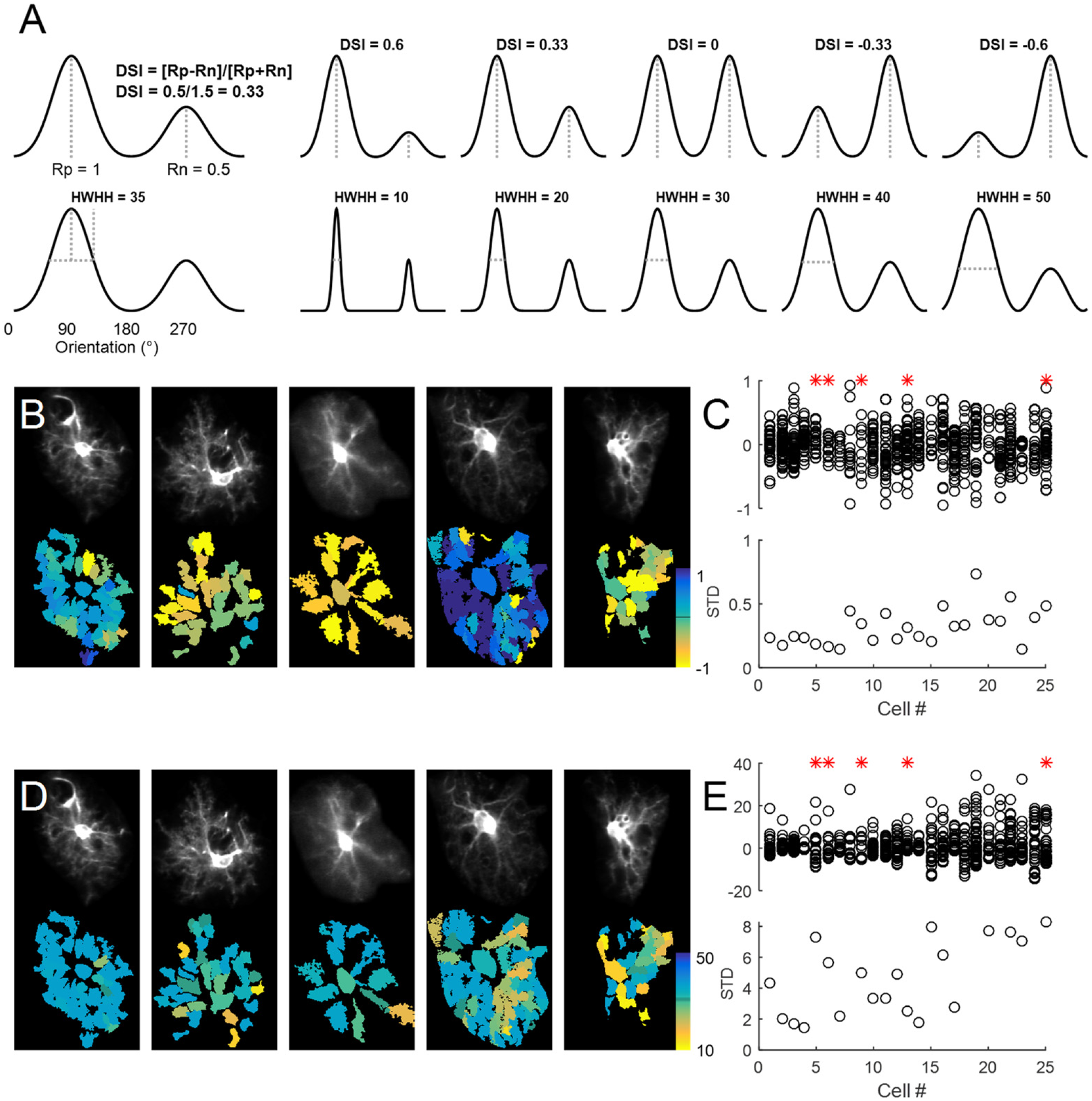
Diversity of other tuning curve parameters. **A.** Schematic depiction of tuning curve parameters. Direction Selectivity Index (DSI) measures the relative response amplitudes to the two opposite directions of motion of the same orientation (180 degrees apart). The Half Width at Half Height (HWHH) measures the width of the tuning curve. **B.** Segment maps of direction selectivity index (DSI) for five example cells. **C.** Scatter plot of difference of segment DSIs (relative to the soma; top) and the STD of segment DSIs for all 25 cells. The cells are sorted by the cSTD of the PO (see Fig 3E). Example cells are indicated by red asterisks. **D.** Segment maps of half width at half height (HWHH) for five example cells. **E.** Scatter plot of difference of segment HWHHs (relative to the soma; top) and the STD of segment HWHHs for all 25 cells. The cells are sorted by the cSTD of the PO (see Fig 3E). Example cells are indicated by red asterisks.

**Figure S6.**
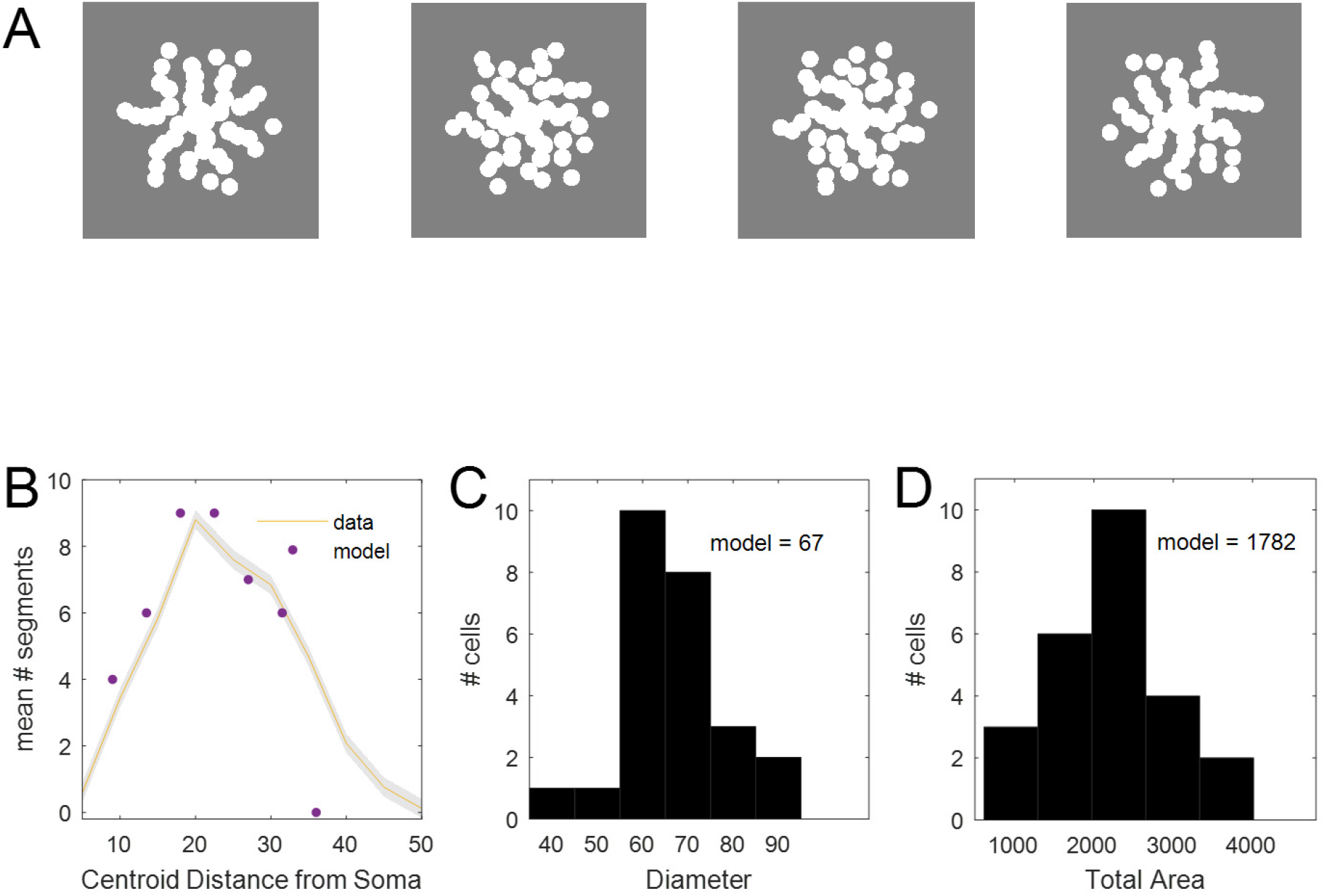
Model astrocyte parameters. **A.** Four example astrocytes generated by the model. The image size is 100μm. **B.** Histogram of the mean (+/- sem) number of segments at each distance from the soma, taken from the output of the ICA from the imaged astrocytes. The dots represent the values used in the model. **C.** Histogram of diameter of the territories from the ICA segments. **D.** Histogram of the total area covered by all the segments.

